# Presence of Newcastle Disease in Vaccinated Indigenous Chicken from Farms in Selected Regions in Kenya- a Cross Sectional Study

**DOI:** 10.1101/2020.07.20.211672

**Authors:** Auleria A. Apopo, Jane M. Ngaira, Jacqueline K. Lichoti, Henry Athiany, Leonard O. Ateya

## Abstract

Indigenous chicken farming is an important source of livelihood for rural families in Kenya. However, the farmers experience challenges including infections from poultry diseases such as Newcastle disease. Vaccination has been used over the years to provide immunity in flocks against disease outbreaks. However, Newcastle diseases virus (NDV) outbreaks are still reported among vaccinated flocks. This study examined the presence of Newcastle disease virus among vaccinated indigenous chicken (IC), in relation to weather (temperature, rainfall, humidity, wind speed), production system, and interspecies interaction. Samples were collected from flocks of indigenous chicken vaccinated with NDV vaccine from 68 households in six counties in Kenya (Bomet, Baringo, Kilifi, Nakuru, Kakamega, and Machakos). Some of the households (n=5 (9%) reported previous ND outbreaks. All the households had other mixed species of birds (ducks, geese, turkey, wild birds). The total number of samples collected was n=1210, oropharyngeal swabs-650, cloacal swab-650. The total number of the pooled sample was 246 pools. The samples were analysed by RT-qPCR targeting the NDV matrix gene. NDV matrix gene was detected in the pooled samples n=177(72%). Out of the positive samples, n=56(32%) were from vaccinated flocks, 91(51%) were from the cloacal swab and 86(49%) from the oropharyngeal swab. The samples were collected during varying weather (temperature, rainfall, humidity and wind speed). There was a statistical significance on the relationship between the presence of NDV and the Vaccination history (p=0.034); production system, (p=0.004), and month of sample collection (p<0.0001). However, there was no significance on the relationship between the presence of NDV and the interaction of IC and other birds. Failure to vaccinate IC results in the presence of the NDV. The free-range production system can have many cases of NDV due to the lack of biosecurity measures. Therefore, farmers should be advised to vaccinate their IC.

## Introduction

Poultry farming accounts for about 30% of agricultural activities. The chicken population in Kenya is approximately 30 million with majority kept as free ranging Indigenous Chicken (IC)^1^. Indigenous chicken is an important component of rural household as a source of food, income, nutrition, insurance against emergencies and has the potential to reduce poverty. Rearing of IC is also linked to different ceremonial roles among communities in Kenya^2^.

Indigenous chicken farming is constrained by several challenges including slow growth rate and maturity rate, poor feeding, high feed cost and high chick’s mortality due to poultry disease. Newcastle disease (ND) is one of the most important infectious diseases of poultry globally and a big constraint in poultry production in rural areas ^3,4^.

Newcastle disease virus (NDV), is the causative agent for the ND ^5^, a single-stranded RNA virus of the genus avulavirus ^6^. NDV is also called Avian Paramyxovirus Virus-1 (APMV-1). APMV-1 virions are pleomorphic in shape, and consist of single-stranded, non-segmented, negative-sense RNA genomes ^7^. The RNA genome contains six major genes that encode the structural proteins which include nucleocapsid (N), phosphoprotein (P), matrix (M), fusion (F), hemagglutinin-neuraminidase (HN) and the RNA dependent RNA (large) polymerase ^7,8^.

In Kenya, different local communities have adopted names for ND depending on the prevailing weather condition when a suspected outbreak is reported such as “muyekha” in western region “Kidere” at the coastal lines, “Kihuruto” at the central area, “Amalda” in the south rift, “Chepkinuch” in north rift and “Mavui” in eastern Kenya. The endemicity of the virus has been attributed to a number of factors; weather, agroecological zoning, disposal of infected carcasses, vaccination processes, interspecies farming and restocking of birds from markets ^9,10^. Prevalence of ND by sero-surveillance in parts of Kenya has been reported to be higher in dry hot zones (17.8%) than cool wet zones (9.9%) and cold seasons^9^. Vaccination has been used to control the disease.

In Kenya, the vaccine used is either live attenuated viral vaccine prepared from “F” strain, live attenuated New Castle disease viral vaccine prepared from the La Sota strain, 1-2 thermostable ND and B strain vaccine. However, there have been reported cases of vaccination failures among the bird population^11^, which has been attributed to several factors including virus shedding, environmental contamination after the previous outbreak, improper application, vaccine neutralization and paternal effect on passive immunity^11,12^. In an experiment to evaluate the effect of vaccination on transmission of highly virulent vaccine^13^, virus shedding was observed in most vaccinated challenged birds compared to those receiving booster vaccination. Other species of birds that are reared alongside chicken, such as ducks, are known to harbor strains of APMV-1 and therefore can transmit the virus to noninfected birds ^3^.

RT-qPCR has been used to detect the different genes of the virus. Detection of the M gene has been done by the use of RT-qPCR for clinical samples by the use of primers from the conserved matrix gene^14,15^. This has been used to detect the lentogenic (low virulence), Mesogenic (moderate virulence) and velogenic (highly virulent) strains of APMV-1^16,4^.

This study reports detection of NDV among vaccinated IC in Kenya detected by RT-qPCR. The study also establishes risk factors, weather (temperature, rainfall, humidity, wind speed), interspecies interaction, Production system of the IC as possible causes of presence of NDV in indigenous chicken.

## Materials and Methods

### Sampling sites and Sample collection

The samples are part of an ongoing targeted ND surveillance done in the country in the year 2017-2018. The samples were collected from six counties in Kenya (Figure 1), during the months of May 2017, June 2017, September 2017 and March 2018, within two agroecological zones (AEZ) III and V. The areas are key in indigenous chicken (IC) farming, trade and cultural significance ^2^. Kakamega, Bomet Nakuru, and Kilifi are within AEZ III, whereas Baringo is within AEZ V. The sampling sites were selected within the sub-counties under the guidance of the regional veterinary government offices. The samples were collected after getting a verbal consent from farmers. Written consent is not a requirement for sample collection in Kenya (KALRO/BIO/KAB/42/36).

**Figure 1:**
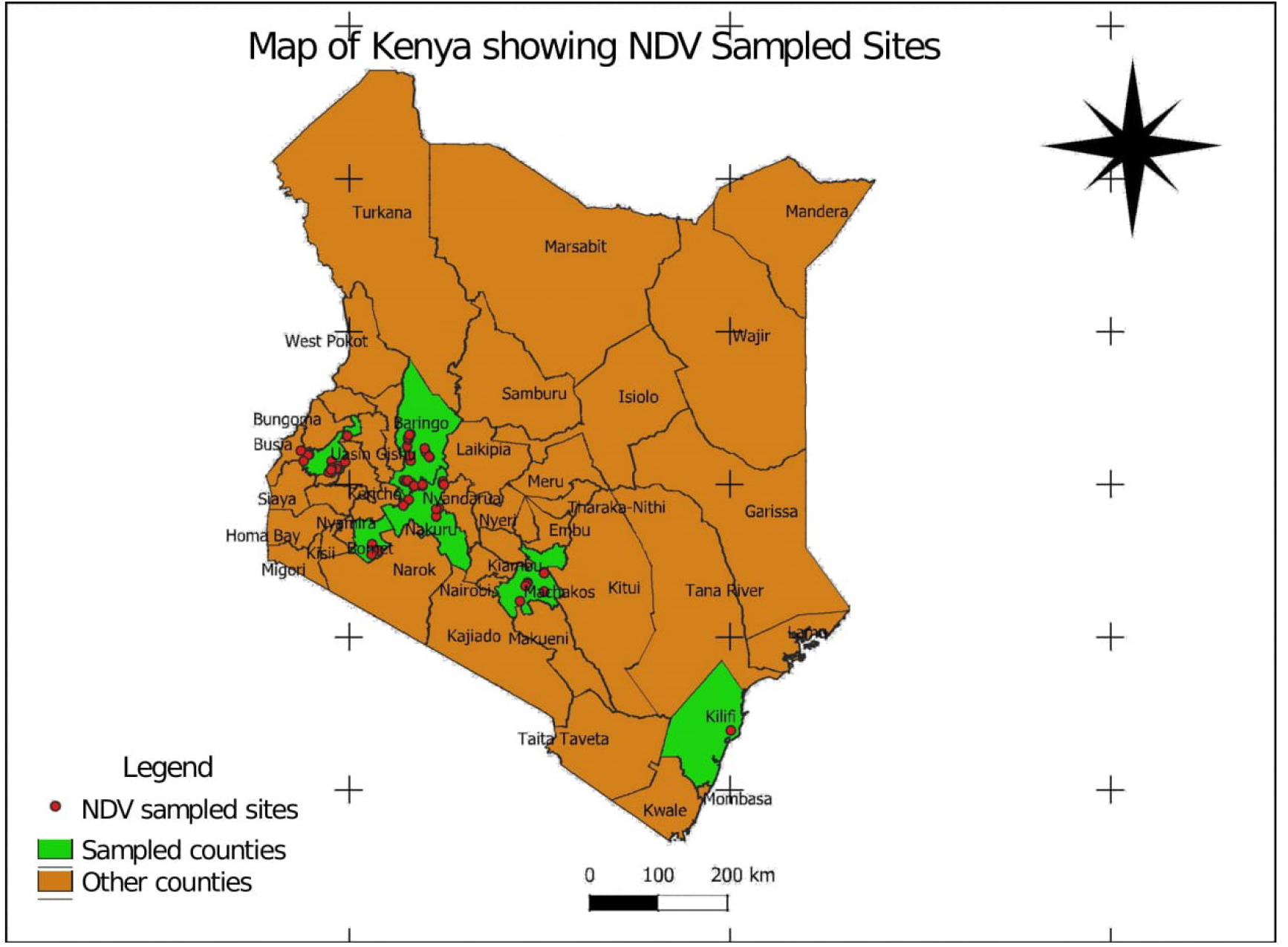
Map of Kenya showing sample collection sites for NDV. Developed using QGIS-3^®^.

The distance from one household to the next was over 1 km. The distance was to ensure that samples in a given region was from unrelated households in terms of interaction of their IC. The total number of birds sampled within a given village/ward was dependent on the total number of samples to be collected in the region. The production systems in homesteads were either the free-range chicken or under a semi-free-range production system. Samples were collected from farms that the IC had either interacted with other mixed species of poultry or not. Before sample collection, verbal consent was sought from the farmer. Data collected from the household included, previous ND outbreaks and vaccination history; source of the birds was collated using the laboratory card. Samples were collected from live domestic indigenous chicken of both sexes and all ages from 3 weeks old. The samples were collected from both healthy birds and birds that symptoms similar to NDV infection. Disposable sterile swabs (polyester or rayon) were used to collect oropharyngeal swab (OP) and cloacal swabs (CL)^17^. The swabs were immediately placed into appropriately labelled cryovials containing 1ml of viral transport media. Samples were then immediately placed in a cool box and later on placed in liquid nitrogen and transported to the labs where they were stored at −80°C awaiting processing ^18^.

### Sample size

The sample size for this study was 1210 samples of both oropharyngeal swabs (650) and cloacal swabs (650) samples from farms across the country. The sample size was calculated using the formula below and it was inclusive of 10% non-response cases

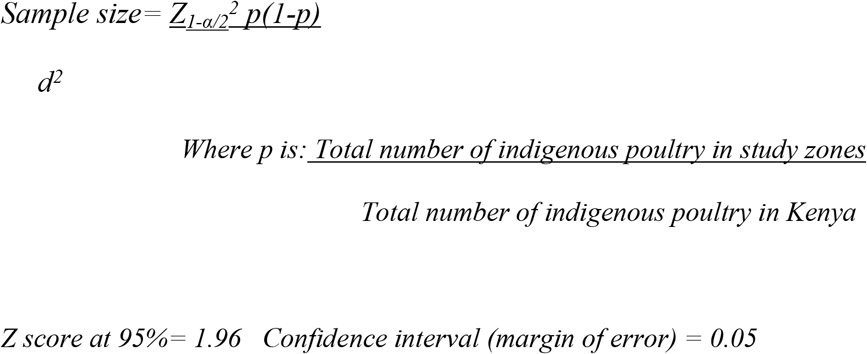

The samples were collected from both farms that had not vaccinated their IC and those that had vaccinated their IC against NDV.

### Sample preparation

Sample processing was done at the Biotechnology Research institute of the Kenya Agricultural Livestock Research organization (KALRO) in accordance with the regulations of the Care and Use of animals Committee from KALRO (KALRO/BIO/KAB/42/36). The samples were pooled based on the region, the household and the type of sample resulting to a total of 246 pooled samples (~5 samples per pool). Pooling was done separately for oropharyngeal swab sample and for cloacal swab sample. Pooling was done by centrifuging the cryovials containing the swabs at 1000xg for about 10 minutes in a refrigerated centrifuge (4°C). This was done to release the virus from the swab to the Viral transport medium. After gently shaking the vial, 100μl of the sample from the same sample collection site and type of sample, were pooled into a sterile cryovial ^17^.

### Virus isolation from eggs

Viral isolation was done using the allantoic route of 9 days old embryonated specific antibody negative (SAN) eggs. The eggs were obtained at day 8 from Kenchic® hatcheries-Nairobi. They were allowed to settle at 37°C. The pooled samples were vortexed at low speed to mix the sample. The eggs were surface disinfected by use of iodine solution before puncturing the egg shell. A volume of 0.2 ml from the pooled sample was inoculated into the allantoic cavity of each of 5 labelled embryonated SAN 9 day old. After inoculation, the eggs were incubated at 35–37°C for 5 days. The eggs were candled daily and the dead were chilled at 4°C awaiting harvesting. The allantoic fluid was harvested aseptically using 2ml syringes into sterile vials. They were then stored at −80°C ^18^.

### RNA extraction and amplification by RT-qPCR

Total RNA was extracted from the allantoic fluid using TRIzol™method-LS (Invitrogen, Carlsbad, US) according to the manufacturer’s instructions. The extracted RNA was amplified by one step RT-QPCR using matrix gene primers. M+4100 (Forward Primer) 5’_-AGTGATGTGCTCGGACCTTC-3’, M+4169 (Matrix Probe) 5’-[FAM]TTCTCTAGCAGTGGGACAGCCTGC[TAMRA]-3’ M-4220 (Reverse Primer) 5’-CCTGAGGAGAGGCATTTGCTA-3’. A Known positive (Lasota Vaccine strain) used as a positive control. The cycling conditions were APMV-1 matrix 40 cycles denaturation 94°C for 10 seconds, annealing 56°C for 30 seconds and extension 72°C 10 seconds ^14^.

### Data analysis

The data was collated using Microsoft excel® 2016. Data analysis was done in SPSS® 2019. CHI square test and Fisher exact test was used to test statistical significance with a p-value 0.05. The analysis was done to test the differences in the relationship between the presence of NDV in vaccinated flocks and other variables (the vaccination history of the IC; interaction with other birds; sample type; production system and Months of sample collection in relation to weather patterns).

## Results

### Samples collected and household vaccination history

Vaccination in this context means that the farmer, through their intervention or animal service provider, had administered the NDV vaccine within the three months inclusive of the sample collection date. The Vaccine administration was orally. The most used vaccine was the Lasota vaccine. From the total number of households, 17 (25%) of these households had vaccinated their indigenous poultry. The vaccination history per county (Figure 2) indicates that all the counties had a higher number IC that were not vaccinated as compared to the vaccinated IC. Some of the households, 5(9%) had previous outbreaks of ND before sample collection. Samples were collected from one household in Kilifi (Sample ID NDVKE-0303-3-F Bird # 17-51) that had an active case of IC having signs and symptoms of suspected NDV. One household in Kakamega (Sample ID NDVKE-0517-37-F Bird # 21-25) had IC with symptoms similar to fowl pox disease. None of the households from Bomet County had vaccinated their IC.

**Figure 2.**
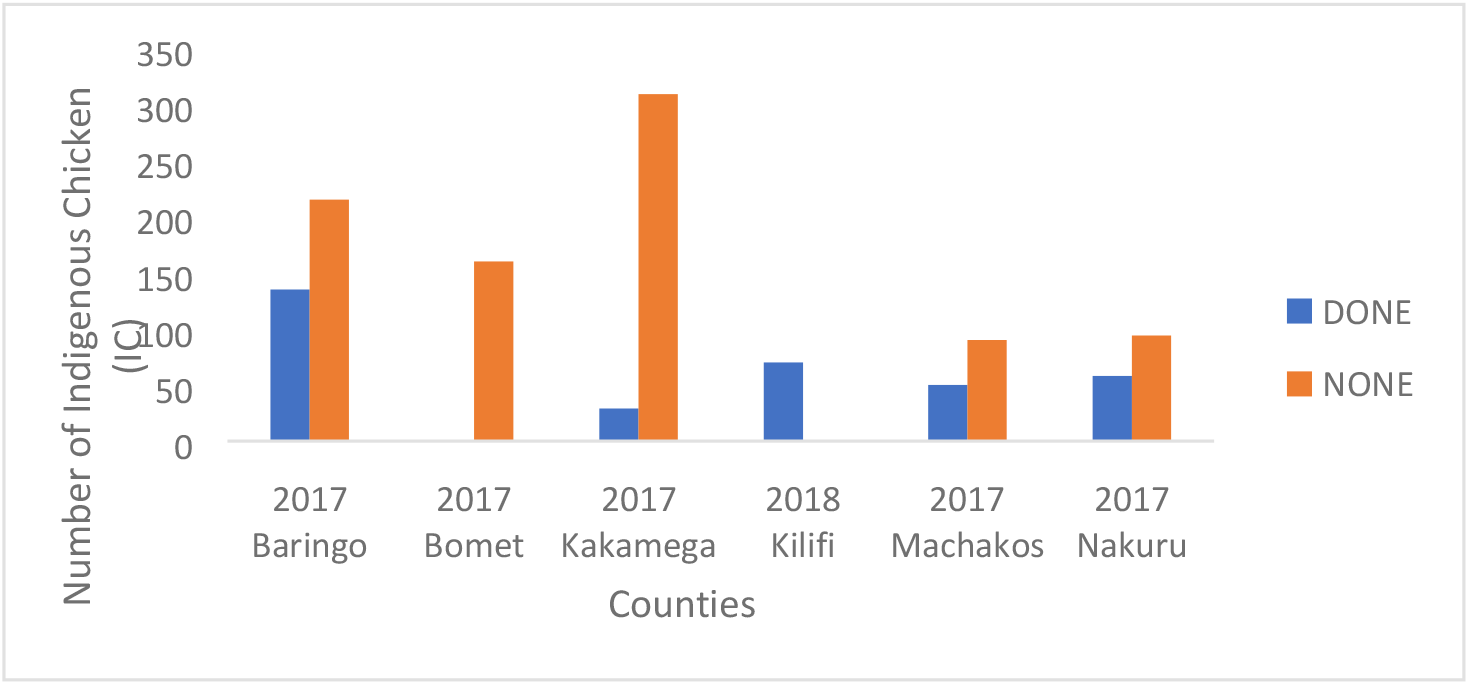
NDV vaccination history per county

### Analysis of RT-qPCR M results

The sample size run through RT-qPCR were 1210 OP (650) and CL (650). The Ct values were an average of 26 to 32. NDV matrix gene was detected in the pooled samples n=177(72%). Out of the positive samples, n= 56(32%) were from vaccinated flocks, 91(51%) were from cloacal swab and 86(49%) from oropharyngeal swabs. The distribution of results for the counties were as follows: Bomet n=25(78.93%), Baringo n=48(66.7%), Kakamega n=44(64.7%), Kilifi n=11(78.5%), Machakos n=25(83.3%), and Nakuru n=24(80%) (Table 1)

**Table 1.**
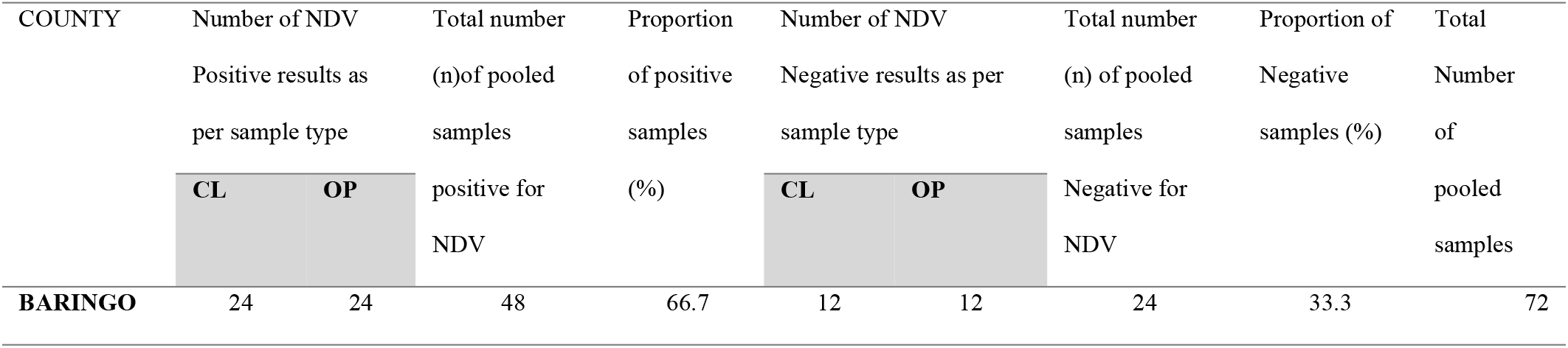
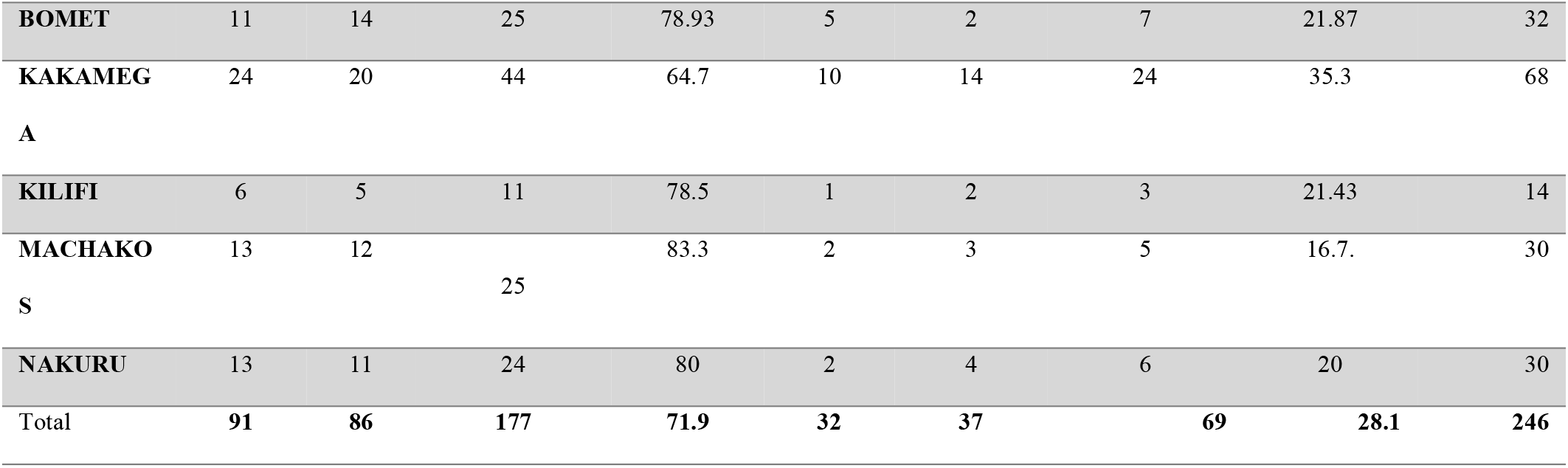
Proportion of positive and negative NDV results from the pooled samples and sample type per County.

**Table 2.**
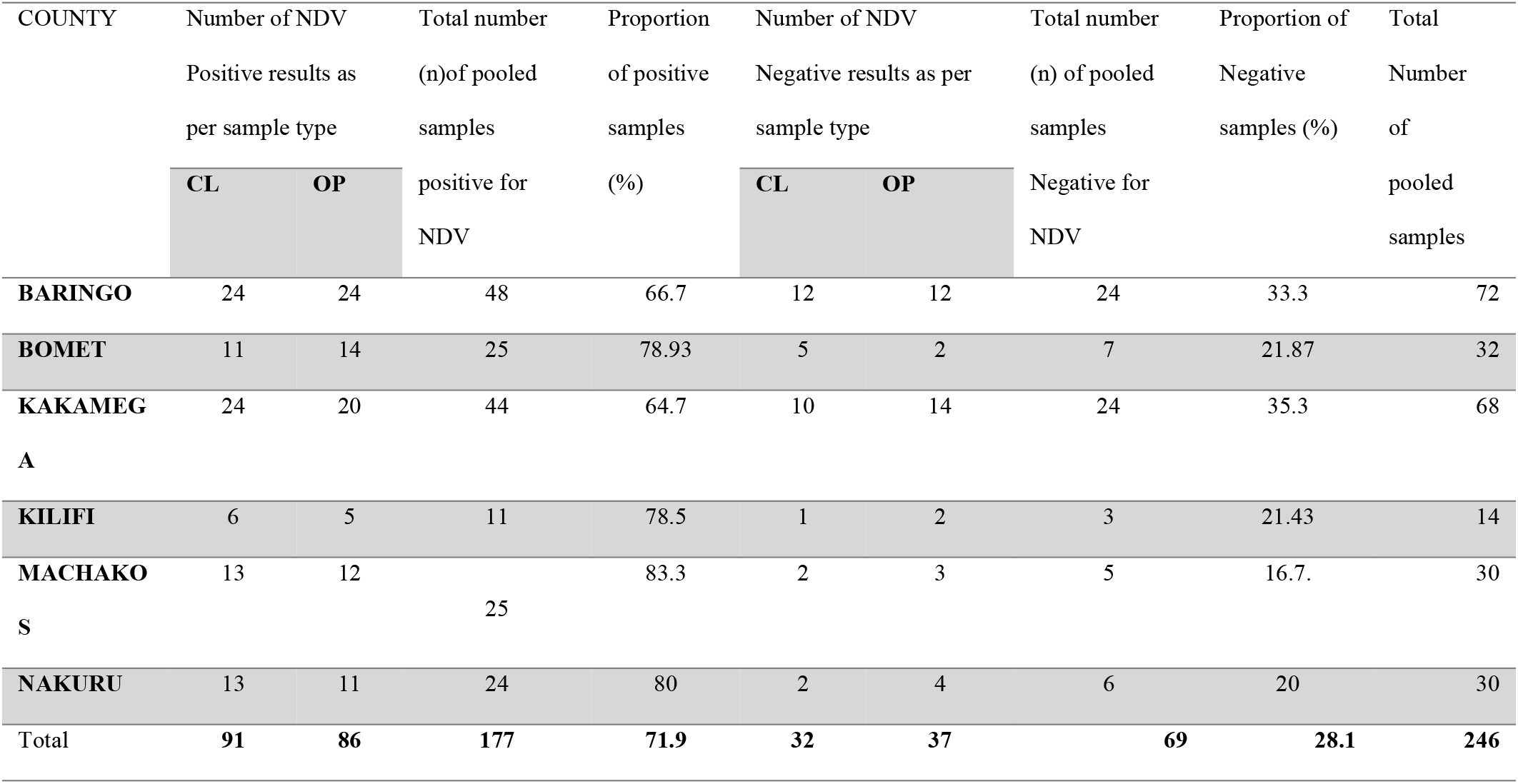
Proportion of positive and negative NDV results from the pooled samples and sample type per County.

### Analysis of the production system

The samples were collected from farms with either of the two production systems; Free range and semi-free range. Most farms kept their IC in the free-range system n=870 (71.9%) (Figure 3). Most of the unvaccinated birds were from the free-range production system as compared to the semi free range system (Figure 4). There was a statistical significance χ^2^ (1, N = 1,210) = 8.458, p=0.004 on the relationship between production system used in the rearing of the IC and the presence of NDV.

**Figure 3.**
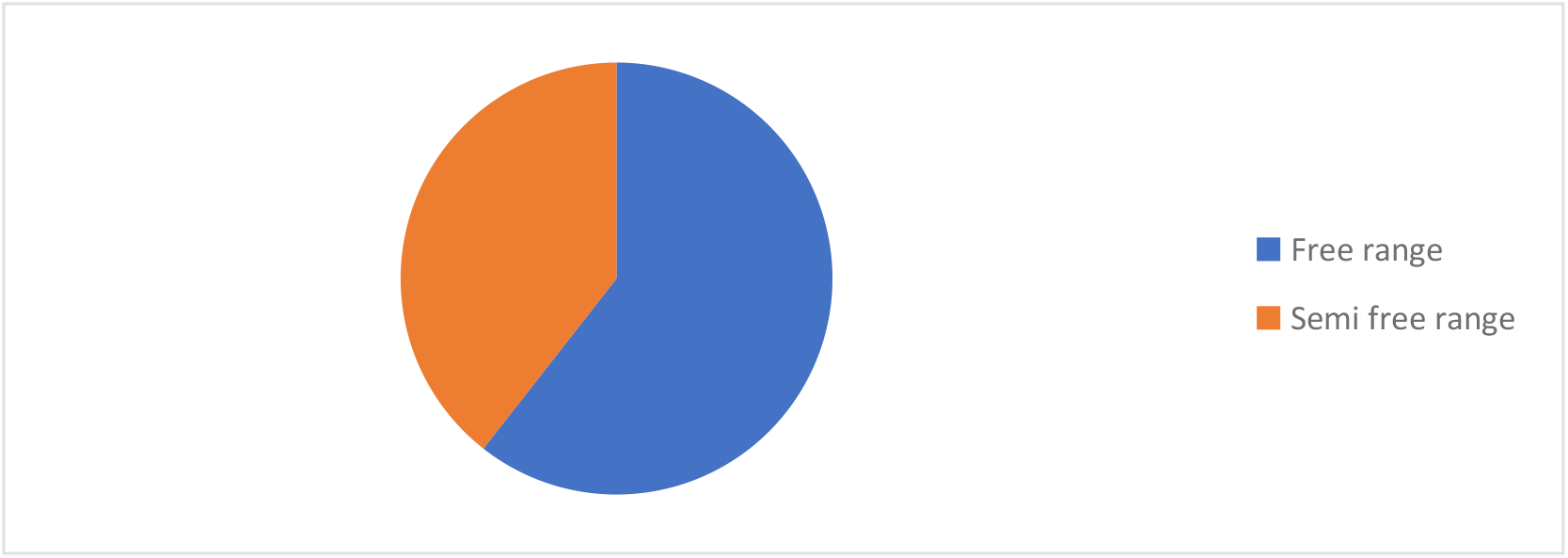
Indigenous production systems in Kenya

**Figure 4.**
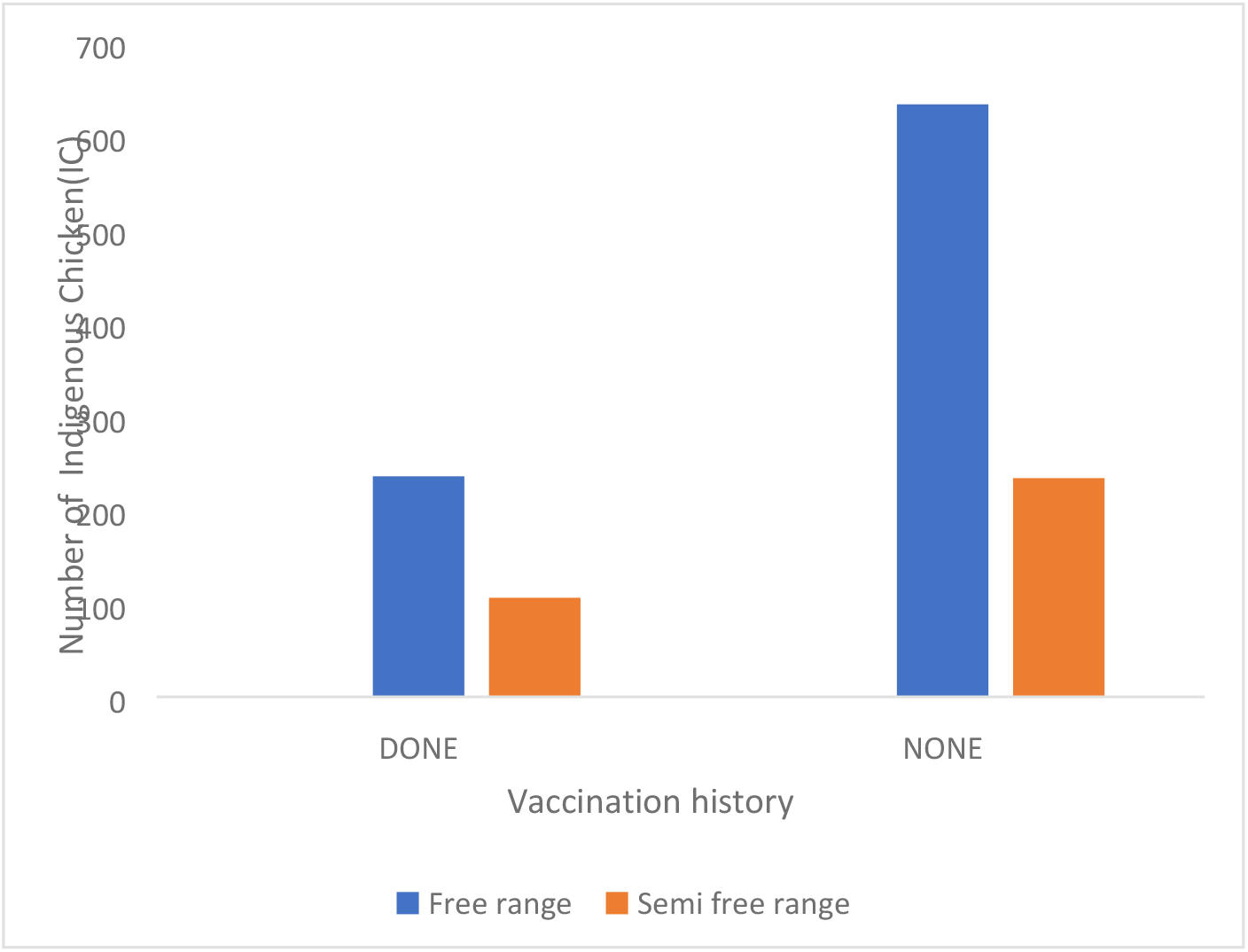
Comparative chart of indigenous chicken production system and the vaccination history in Kenya.

### Analysis of the IC vaccination history and PCR results

The Positive NDV was detected more in IC that were not vaccinated than those vaccinated. (Figure 4) There was a statistical significance χ^2^ (1, N = 1,210) = 4.051, *p=0.034* on the relationship between Vaccination history of the IC and the presence of NDV.

**Figure 5.**
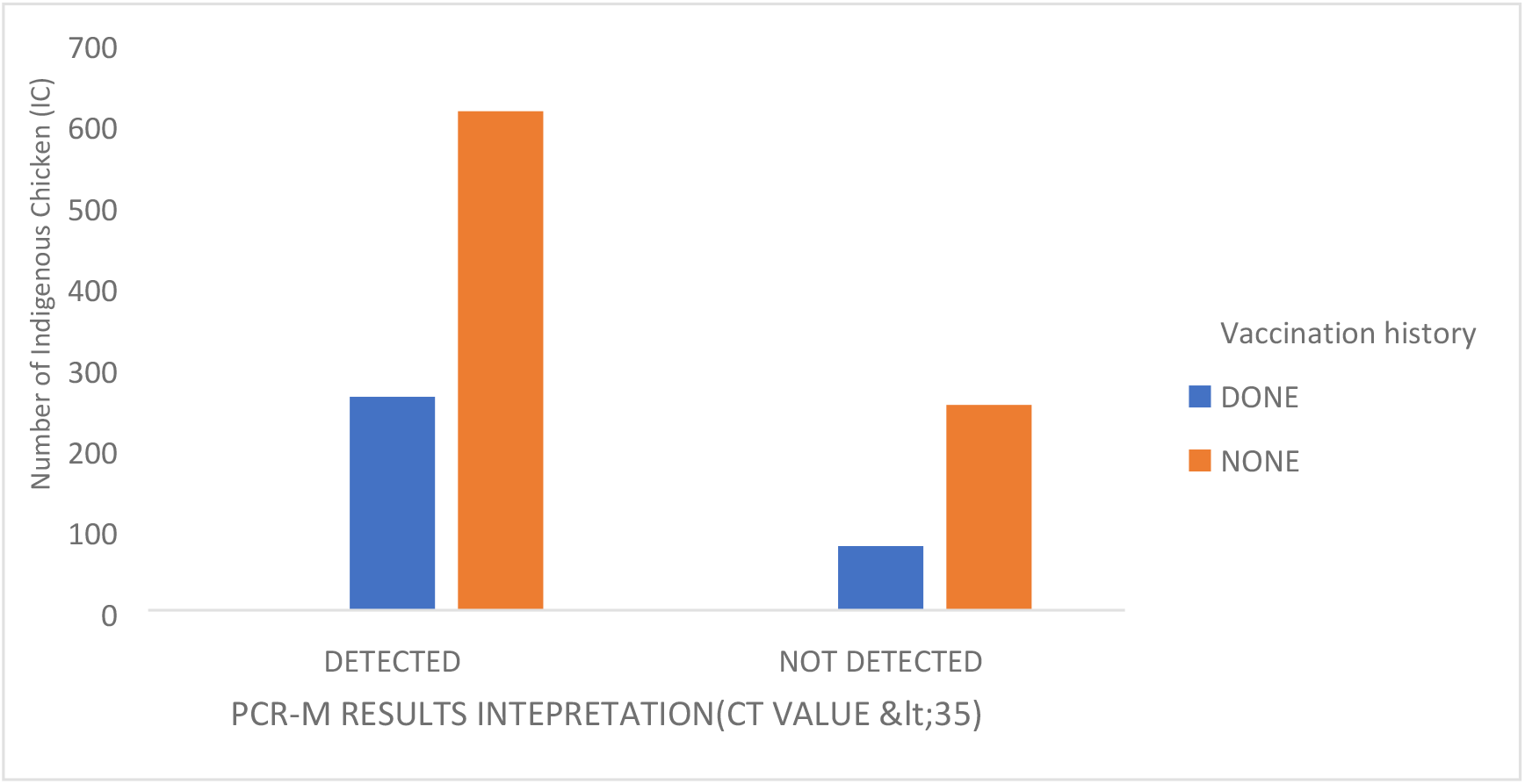
Comparative chart on the presence of NDV and the vaccination history in Kenya

### Comparison of RT-qPCR results with interaction with other birds

The analysis was done to compare the relationship between the presence of NDV interacting with other domestic birds such as Turkey, ducks and goose. There was no statistical significance χ^2^ (1, N = 1,210) = 2.027, p=0.155 on the relationship between interaction with other birds and the presence of NDV.

### Comparison of RT-qPCR results with month and prevailing weather condition

Analysis of the relationship between PCR results and month of sample collection showed a statistical significance χ^2^ (3, N = 1,210) = 24.327, p<0.0001. (Figure 7)

**Figure 6.**
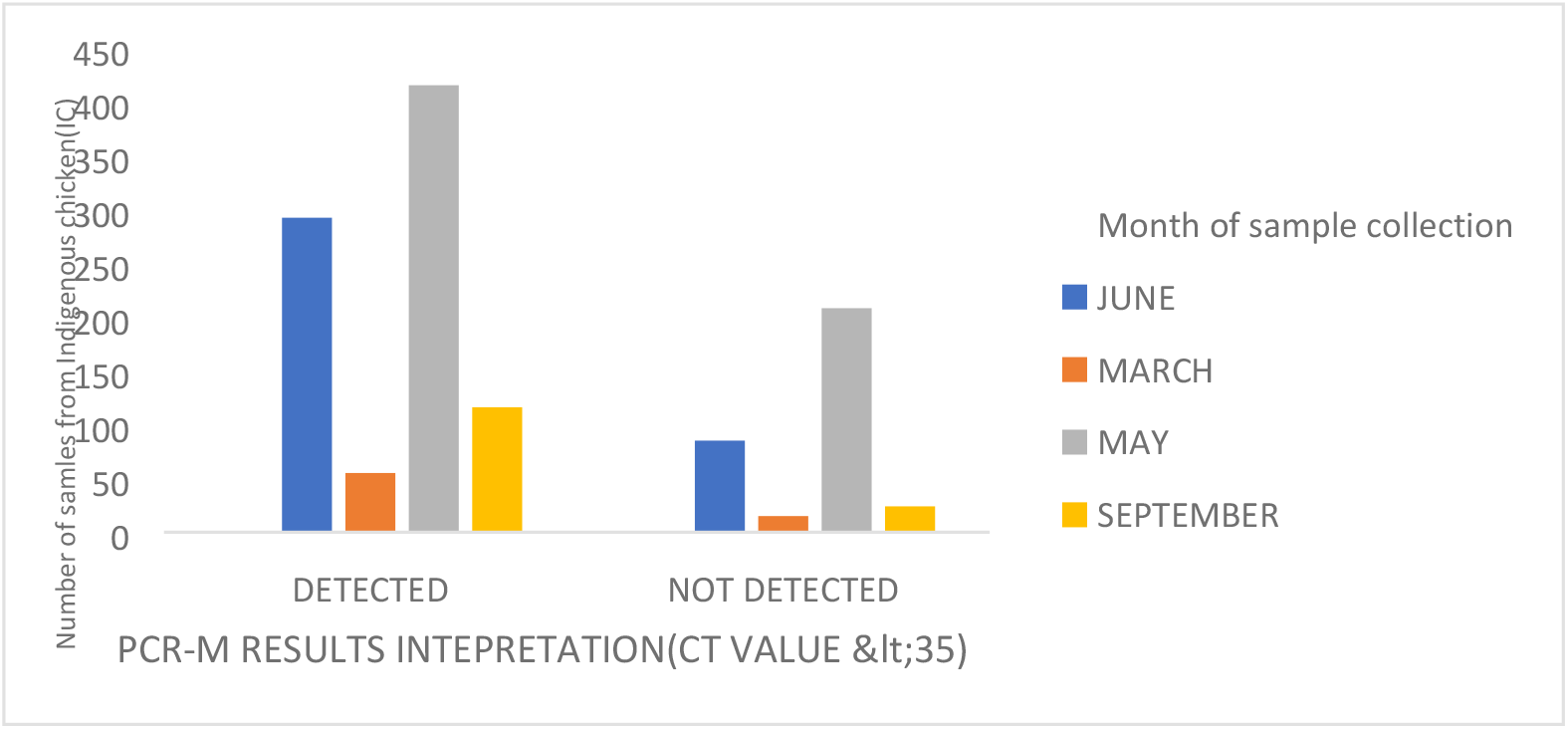
Detection of NDV from samples collected at different months in Kenya

**Figure 7:**
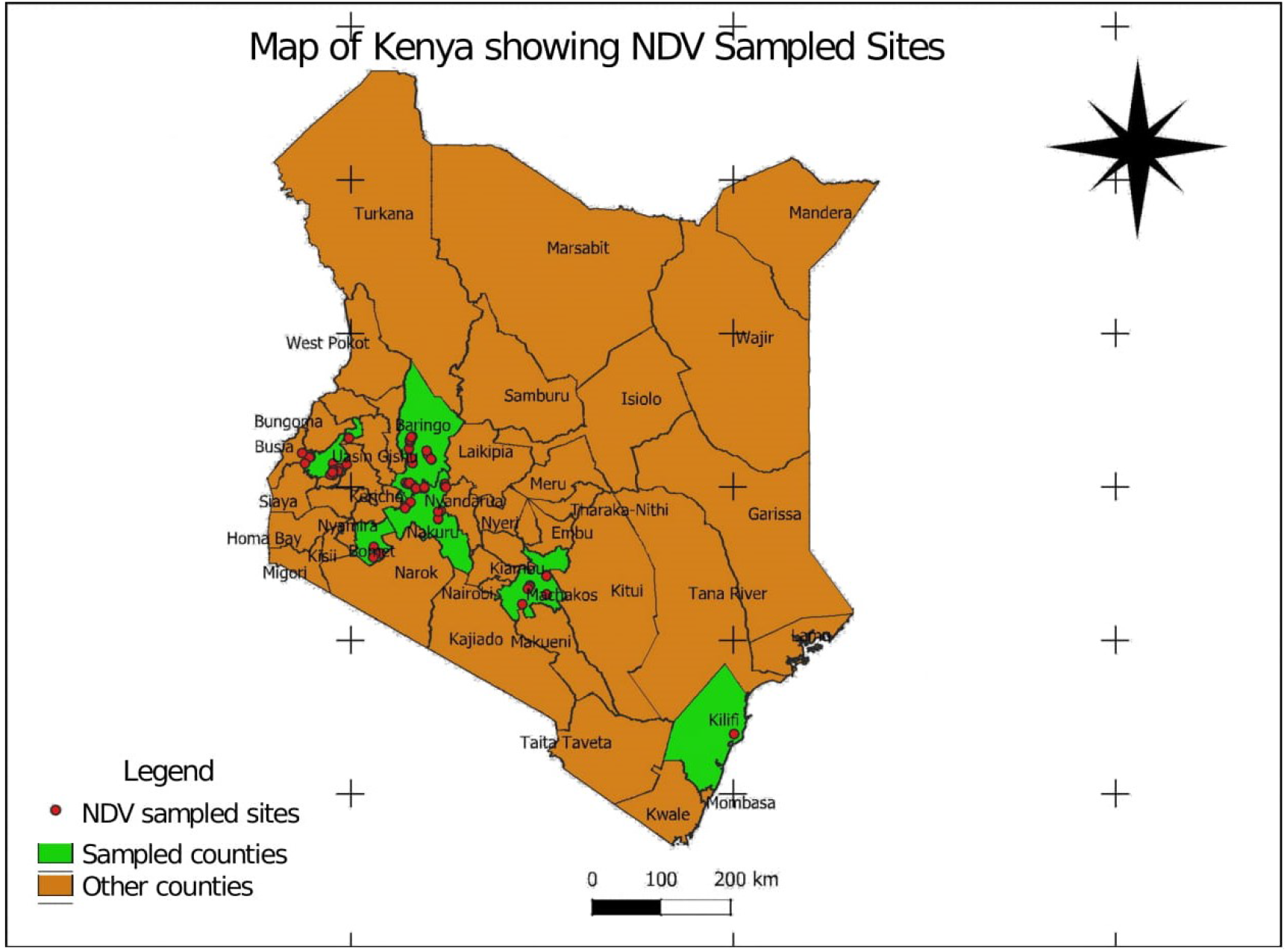
Map of Kenya showing sample collection sites for NDV.Developed using QGIS-3®.

**Figure 8.**
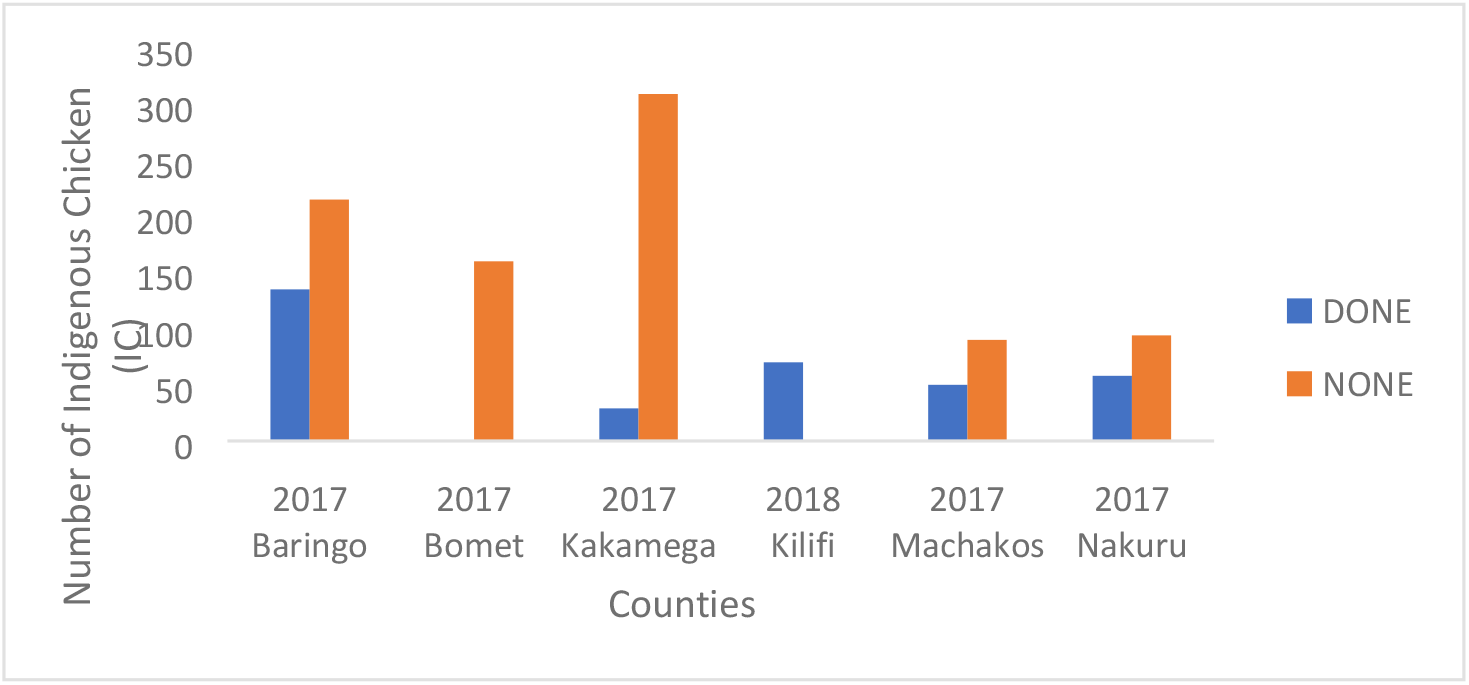
NDV vaccination history per county in Kenya

**Figure 9.**
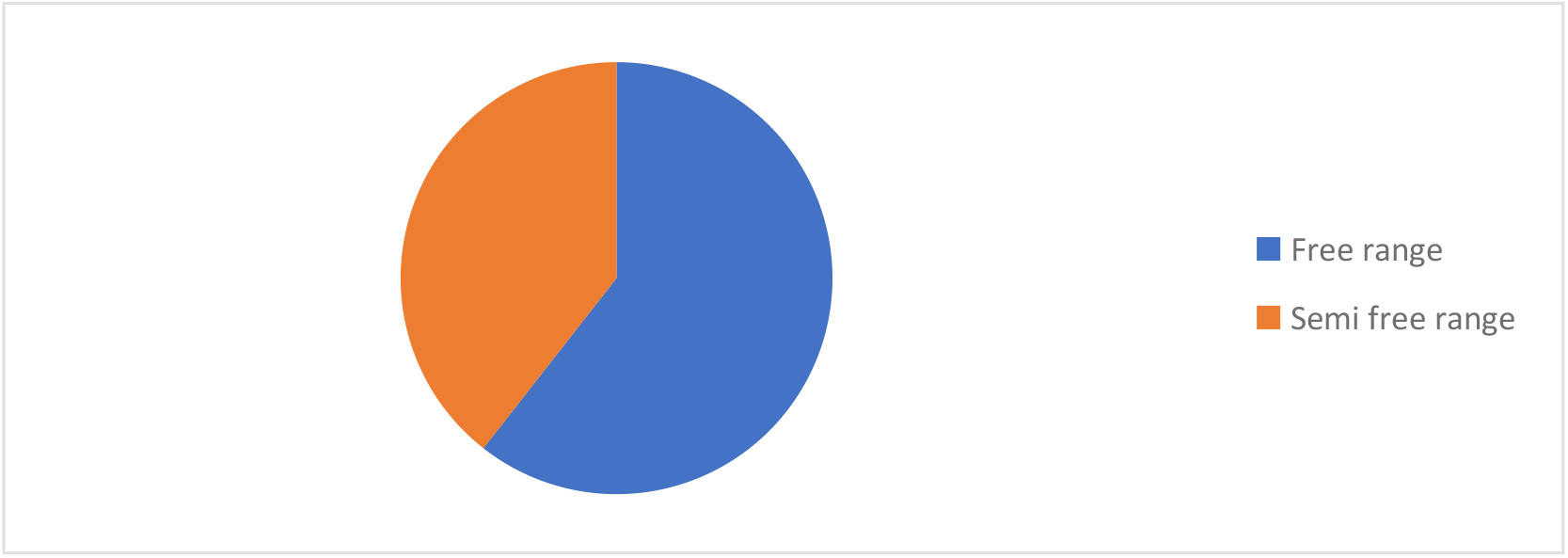
Indigenous production systems in Kenya

**Figure 10.**
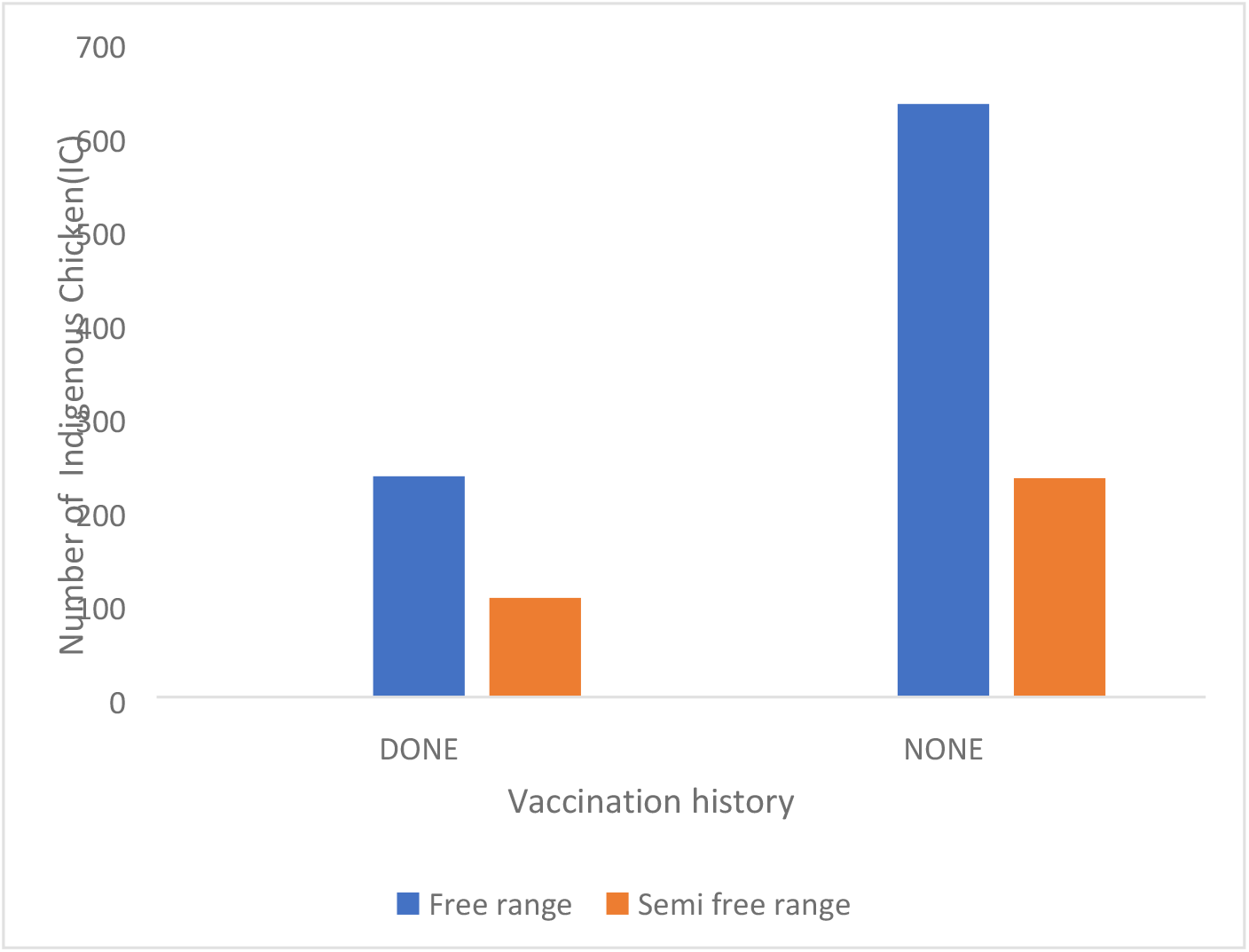
Comparative chart of indigenous chicken production system and the vaccination history in Kenya.

**Figure 11.**
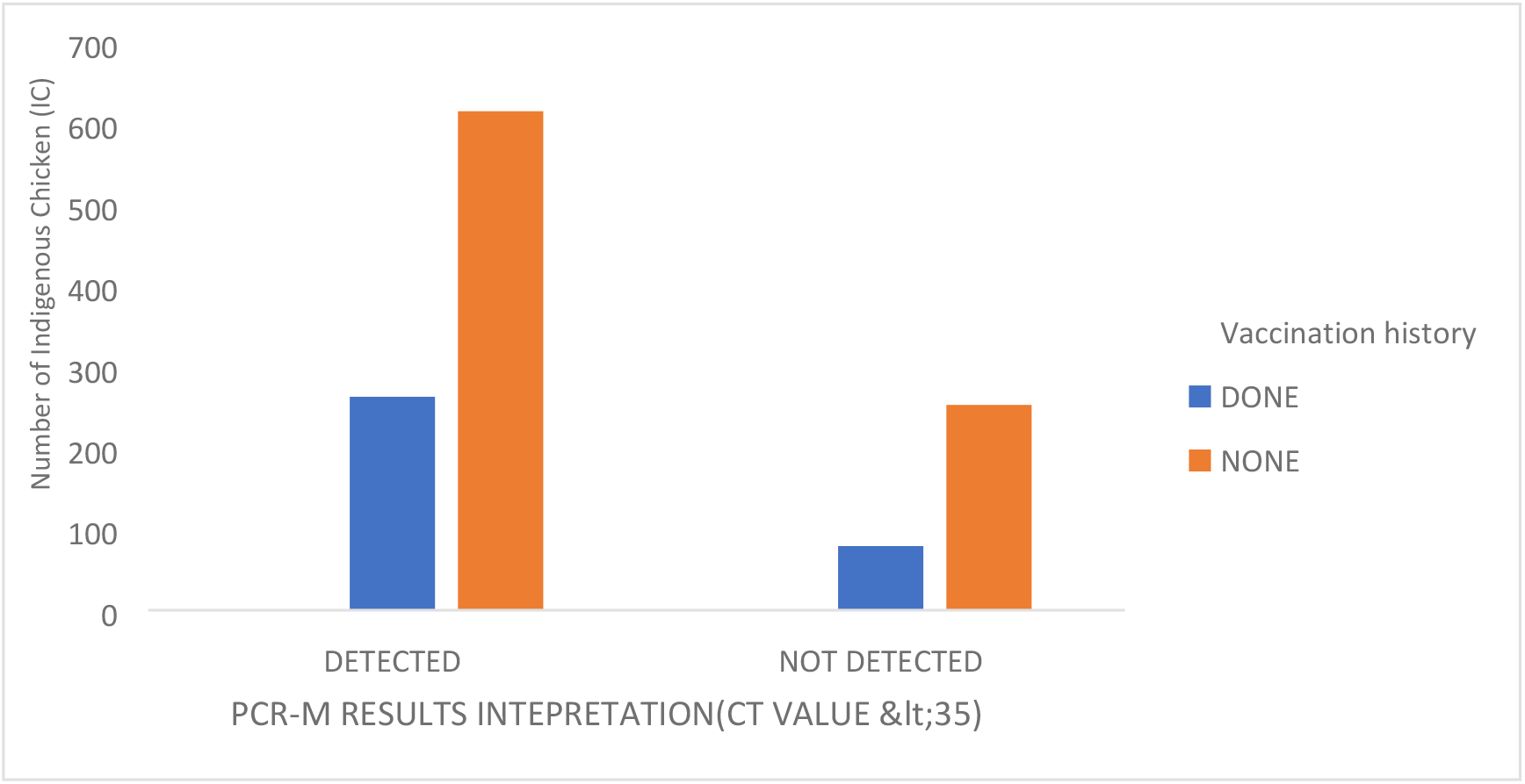
Comparative chart on the presence of NDV and the vaccination history in Kenya

**Figure 12.**
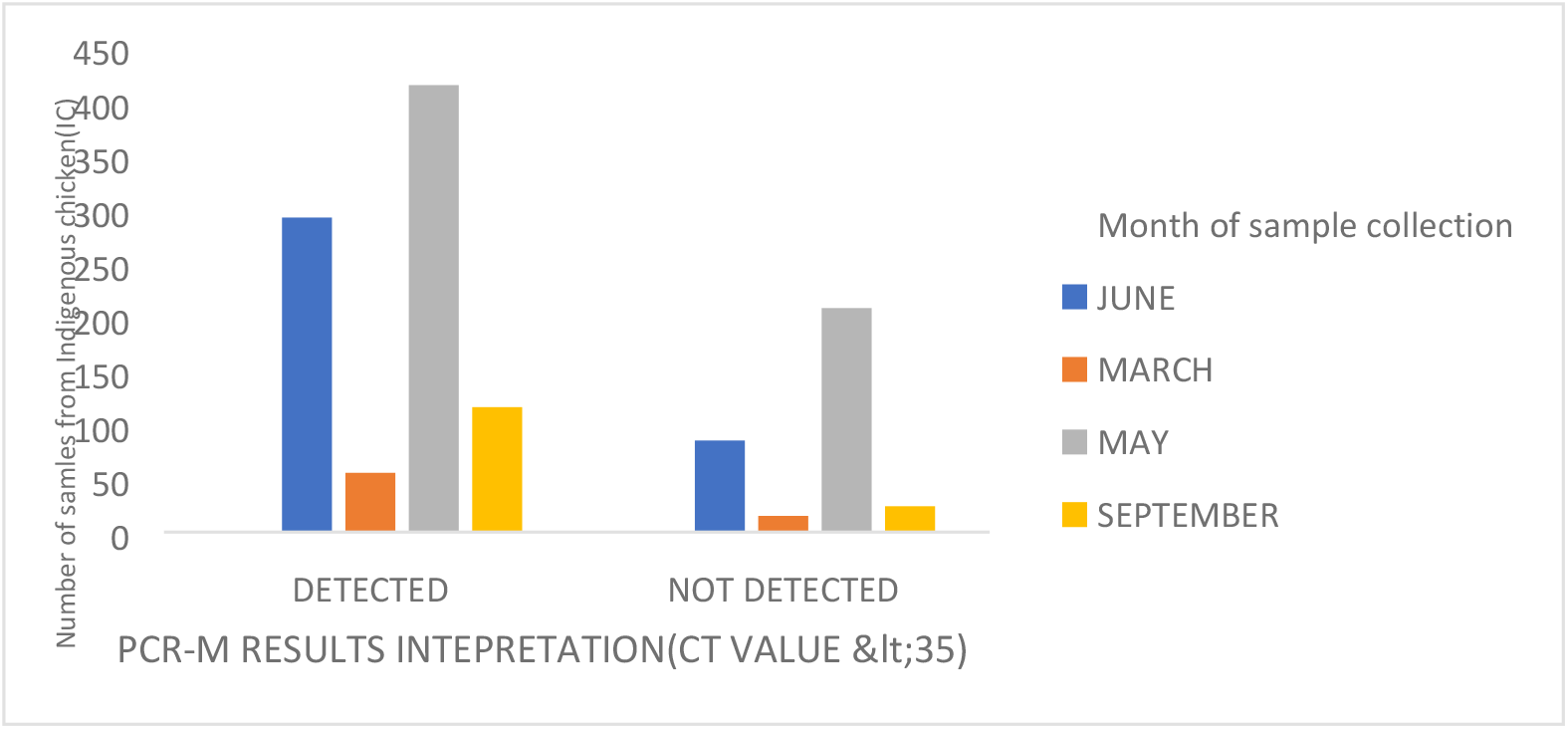
Detection of NDV from samples collected at different months in Kenya

## Discussion

Vaccination of chicken in Kenya is done mainly by use of the Lasota vaccine (Avivax-L) from Kenya Vaccine Production Institute (KEVEVAPI). Vaccination is routinely done on the commercial exotic breeds than the indigenous chicken (IC). In this study, we noted only 17(25%) of the households had vaccinated their IC. It was noted during the sample collection, that IC farmers were aware of Newcastle disease, its clinical sign, its effect on the poultry population and the importance of vaccination. However, the uptake of vaccination was low. The low uptake was due to the small flock size kept by the farmers compared to the package dose of the vaccine, the cost of the vaccine and the lack of appropriate cold chain to transport the vaccine. This can be attributed to the assumption that IC are hardy and do not require vaccination^19–22^. The association of presence of NDV and Vaccination history (p=0.034) indicates that failure to vaccinate IC results in increased NDV cases.

This study results show that NDV cases were detected in some vaccinated flocks (32%). The detection of the matrix gene of the virus is an indication of the presences of the viral antigen in either the respiratory tract (OP) or intestinal tract (CL). The matrix gene detects the conservative gene of the NDV virus of all virulence (Low, moderate and velogenic strains)^23^. Therefore, the presence of the virus can be attributed to viral shedding. This can lead to subsequent transmission of the virus to susceptible flocks^11,13,24^. In Kenya, partial characterization of the ND virus showed that the Lasota vaccine has an 83% similarity with strains circulating in Kenya^25^. The Kenyan genotype is genotype V whereas the Lasota genotype is genotype II of avian paramyxoviruses showing a slight difference in the geneology of the strains.

The production systems for indigenous chicken in the regions we carried out the field study were either free-range and semi free-range systems^25^. There was a significant relationship in the presence of NDV and the production system (p=0.004). In such cases, biosecurity practices are minimal and interaction of chicken from one neighbourhood leads to the rapid spread of ND and high mortalities of birds in households in case of an outbreak. This may contribute to the endemicity of the NDV in Kenya ^2,20,26^.

Prevailing weather conditions such as strong winds, high temperatures, and relatively dry weather results in stress in birds and lowered immunity. Our study reported that the samples were collected during month days that are relatively warm with minimum rainfall. There was a statistical significance between the presence of NDV to the month of sample collection (p<0.0001). Dry weather, therefore, translates to limited access to food, especially for the scavenging birds, stress due to hunger, resulting in lowered immunity and increased cases of NDV^9,27,28^. The farmers were aware that the disease occurred just before the rainy season and attributed this to strong winds, hence the origin of the local names^2^

Traditionally, Indigenous poultry farming in Kenya was reared for its cultural significance^2^.Cultural practices such as gifting have been done over the years. The new birds are introduced into the flock with no history of ND infection. These new birds may transmit the disease through the shedding of the virus if they were previously vaccinated and the new farm had no history of vaccination ^28,11^. In many farms in Kenya cockerels are “borrowed” from neighbours to sire chicks in farms. In this case the cockerel, through paternal inheritance, may genetically influence the ability of offspring to maintain maternal antibody levels against ND ^12^.

Other species of birds, ducks, turkey and feral birds are known to be a source of NDV. Ducks are known to be resistance to the disease, exhibiting no clinical signs and can be a source of shedding of the virus ^3,29^. Therefore, in a mixed-species farm, the birds can harbour the virus which in turn can affect the existing flock, new flock or any immunocompromised flock. In the study, there was no significant association between presence of disease and the interaction of the IC with the other domestic birds.

## Conclusion

ND is an endemic disease in the country and all IC in the areas of study are prone to infection. Vaccination should be used as a way to control the disease. However, the timing of the vaccine with regard to weather conditions should be considered. From the study, the ideal time for vaccination is during the onset of dry season when there is lowered immunity. Sensitization of indigenous poultry farmers on importance of vaccination should done. Sensitization on timing of vaccination and provision of smaller doses of vaccines would go a long way in controlling ND in Kenya.

There is a need to identify the strain of NDV circulating in Kenya by molecular characterization of its complete genome. This would help know the appropriate vaccine for circulating strains minimizing virus shedding.

Identifying strains circulating among wild bird in comparison to strains in domestic birds will help in both developments of new diagnostic kit and improve detection of NDV. Laboratory detection of NDV should include targeting the matrix gene. This should be done so that we don’t fail to detect low virulence NDV that may be transmitted to other IC of lower immunity through shedding.

## Acknowledgement

I acknowledge the Director of Veterinary Services for giving me the chance to undertake this research. Sheila Apopo, Dorcus Omoga and Steven N. for your endless support and advice. The staff at Kakamega, Kilifi, Baringo Nakuru, and Machakos county veterinary offices for your support in the field.

## SUPPLEMENTARY DATA

*Supplementary data-Temperature, rainfall, humidity and wind speed of the different location during sample collection (courtesy Kenya meteorological department.*

| Station/Year/Month | Max temp. | Min temp. | Total rainfall | RH 9am | RH 3PM | Wind run in km/day |
| --- | --- | --- | --- | --- | --- | --- |
| Kakamega May 2017 | 27.9 | 16 | 217.5 | 81% | 65% | 56.2km/day |
| Kilifi (Mtwapa) March 2018 | 31.9 | 24 | 98.3 | 80% | 69% | 80.5km/day |
| Machakos September 2017 | 25.5 | 12.1 | 5.7 | 71% | 43% | 124.2km/day |
| Nakuru June 2017 | 26.6 | 11.1 | 36.7 | 73% | 41% | 104.3km/day |
| Bomet*(May 2017) | 24.4 | 13 | 147 |  |  |  |
| Baringo*(May 2017) | 32.4 | 17.5 | 100 |  |  |  |

\* Data obtained from climate-data.org

